# Content tuning in the medial temporal lobe cortex: Voxels that perceive, retrieve

**DOI:** 10.1101/635128

**Authors:** Heidrun Schultz, Roni Tibon, Karen F. LaRocque, Stephanie A. Gagnon, Anthony D. Wagner, Bernhard P. Staresina

## Abstract

How do we recall vivid details from our past based only on sparse cues? Research suggests that the phenomenological reinstatement of past experiences is accompanied by neural reinstatement of the original percept. This process critically depends on the medial temporal lobe (MTL). Within the MTL, perirhinal cortex (PRC) and parahippocampal cortex (PHC) are thought to support encoding and recall of objects and scenes, respectively, with the hippocampus (HC) serving as a content-independent hub. If the fidelity of recall indeed arises from neural reinstatement of perceptual activity, then successful recall should preferentially draw upon those neural populations within content-sensitive MTL cortex that are tuned to the same content during perception. We tested this hypothesis by having eighteen human participants undergo functional magnetic resonance imaging (fMRI) while they encoded and recalled objects and scenes paired with words. Critically, recall was cued with the words only. While HC distinguished successful from unsuccessful recall of both objects and scenes, PRC and PHC were preferentially engaged during successful vs. unsuccessful object and scene recall, respectively. Importantly, within PRC and PHC, this content-sensitive recall was predicted by content tuning during perception: Across PRC voxels, we observed a positive linear relationship between object tuning during perception and successful object recall, while across PHC voxels, we observed a positive linear relationship between scene tuning during perception and successful scene recall. Our results thus highlight content-based roles of MTL cortical regions for episodic memory and reveal a direct mapping between content-specific tuning during perception and successful recall.

## 1 Introduction

One of the most intriguing features of the human brain is its ability to recall vivid episodes from long-term memory in response to sparse cues. For example, the word ‘breakfast’ may elicit recall of visual information including spatial (e.g. a bright kitchen) and object details (e.g. a croissant). This phenomenological reinstatement of past experiences is mirrored in cortical reinstatement – a neural reactivation of the original perceptual trace (Danker and Anderson, 2010). The medial temporal lobe (MTL) and its subregions play a key role in recall (Zola-Morgan and Squire, 1990; Eichenbaum et al., 2007). Anatomically, the MTL’s input/output regions, perirhinal cortex (PRC) and parahippocampal cortex (PHC), have differentially weighted reciprocal connections to the ventral and dorsal visual stream, respectively (Suzuki and Amaral, 1994a; Lavenex and Amaral, 2000; van Strien et al., 2009). They are therefore well-suited to relay content-sensitive signals from sensory areas to the hippocampus (HC) during perception and encoding and vice versa during retrieval. Indeed, these parallel information streams converge in the HC, enabling it to support memory in a content-independent manner (Davachi, 2006; Eichenbaum et al., 2007; Danker and Anderson, 2010). In support of this view, human functional imaging studies have linked object-related vs. spatial processing to PRC vs. PHC for a range of tasks, including perception (Litman et al., 2009), context encoding (Awipi and Davachi, 2008; Staresina et al., 2011), reactivation after interrupted rehearsal (Schultz et al., 2012), and associative retrieval of object-scene pairs (Staresina et al., 2013b). Conversely, the HC, instead of representing perceptual content, is thought to store indices linking distributed cortical memory traces (Teyler and DiScenna, 1986; Teyler and Rudy, 2007), thereby well-suited to coordinate pattern completion from partial cues (Marr, 1971; Norman and O’Reilly, 2003; Staresina et al., 2012; Horner et al., 2015).

The reciprocity of MTL connectivity implies overlapping activity profiles between perception and retrieval in content-sensitive pathways, and is thought to underlie cortical reinstatement (Eichenbaum et al., 2007; Danker and Anderson, 2010). Indeed, there is evidence that neural activity that was present during the original encoding of a memory is reinstated during retrieval, as demonstrated using univariate analyses of encoding-retrieval overlap (Nyberg et al., 2000; Wheeler et al., 2000; Kahn et al., 2004), correlative encoding-retrieval similarity measures (Staresina et al., 2012; Ritchey et al., 2013), and multivariate decoding approaches (Polyn et al., 2005; Johnson et al., 2009; Mack and Preston, 2016; Liang and Preston, 2017). Moreover, cortical reinstatement scales with the reported fidelity of recall (Kuhl et al., 2011; Kuhl and Chun, 2014). The precise topographical mapping of content-sensitivity at perception to cortical reinstatement at retrieval, however, is unclear. If cortical reinstatement reflects a restoration of a distinct neural state during the original encoding experience, then successful recall of content should predominantly draw on neural populations that distinguished the content from others during perception. That is, the more content-tuned neural populations are during perception, the more diagnostic they should be of successful recall of their preferred content.

Here, we investigated content-sensitivity of MTL subregions during episodic memory recall, and how it maps to content tuning during perception. To this end, we had participants undergo fMRI while they encoded and retrieved adjectives paired with an object or scene image. During retrieval, they only saw the adjective cue and tried to recall the associated object or scene. If HC contributes to recall in a content-independent fashion (as predicted by MTL connectivity), we would expect similar involvement during cued recall of both objects and scenes. Conversely, since MTL anatomy predicts content-sensitivity in PRC and PHC, we expect a preference for object recall in PRC and for scene recall in PHC. Critically, within PRC, we expect a positive correlation such that voxels exhibiting stronger object tuning during perception should be recruited more strongly for successful object recall. In contrast within PHC, we expect a positive correlation such that voxels exhibiting stronger scene tuning during perception should be recruited more strongly for successful scene recall.

## 2 Materials and Methods

### 2.1 Participants

A total of 34 volunteers (all right-handed, native English speakers, normal or corrected-to-normal vision) participated in the fMRI experiment. Sixteen participants were excluded from data analysis. Of those, one was excluded due to excessive movement, and one due to non-compliance. Fourteen datasets suffered data loss due to scanner malfunction. The results of the remaining n=18 participants (11 female; mean age 22.7 yrs, range 18-33 yrs) are reported here. All participants gave written informed consent in a manner approved by the local ethics committee, and were paid for their participation.

### 2.2 Stimuli and Procedure

Stimuli consisted of 60 images of objects and 60 images of scenes (Konkle et al., 2010a, 2010b) as well as 120 English adjectives (Staresina et al., 2011). An additional 5 objects, 5 scenes, and 10 adjectives were used for practice. Per stimulus subcategory (e.g. desk, garden, etc.), only one image was used. Adjective-image pairs were randomized for each participant.

During fMRI, participants viewed stimuli via projection to a mirror mounted on the head coil, and responded using an MR compatible button box. The fMRI task (Figure 1A) used a slow event-related design, consisting of four runs (two object runs, two scene runs). Object and scene runs were presented in an alternating order that was counterbalanced across participants. Each run included an encoding and a retrieval phase (30 trials each), as well as pre- and post-encoding resting phases (3 minutes each). In each trial of the encoding phase, participants saw an object or scene image (400 × 400 pixels) presented in the center of the screen together with an adjective. Participants were asked to press the left or right button on a right-hand button box if they thought the adjective and image matched or did not match, respectively (“decide whether the adjective could be used to describe the image”). Adjective-image pairs were presented for 5s, followed by 10s of an arrows task (active baseline task) (Stark and Squire, 2001) during which participants indicated the direction of left- or right-pointing arrows by pressing the left or right button. In the retrieval phase, the adjectives from the encoding phase were presented again in randomized order. Adjectives were presented for 5s, and participants were asked to press the left button if they successfully recalled the associated image, and the right button if they did not. Each retrieval trial was again followed by 10s of the arrows task. Before and after encoding, participants additionally engaged in an odd-even numbers task for 180s (offline resting phase), separated from the task phases by a transition screen (10s each). In the odd-even task, participants were presented with random numbers between 1 and 99 and pressed the left button for even numbers and the right button for odd numbers. Altogether, each run lasted 22 min.

**Figure 1.**
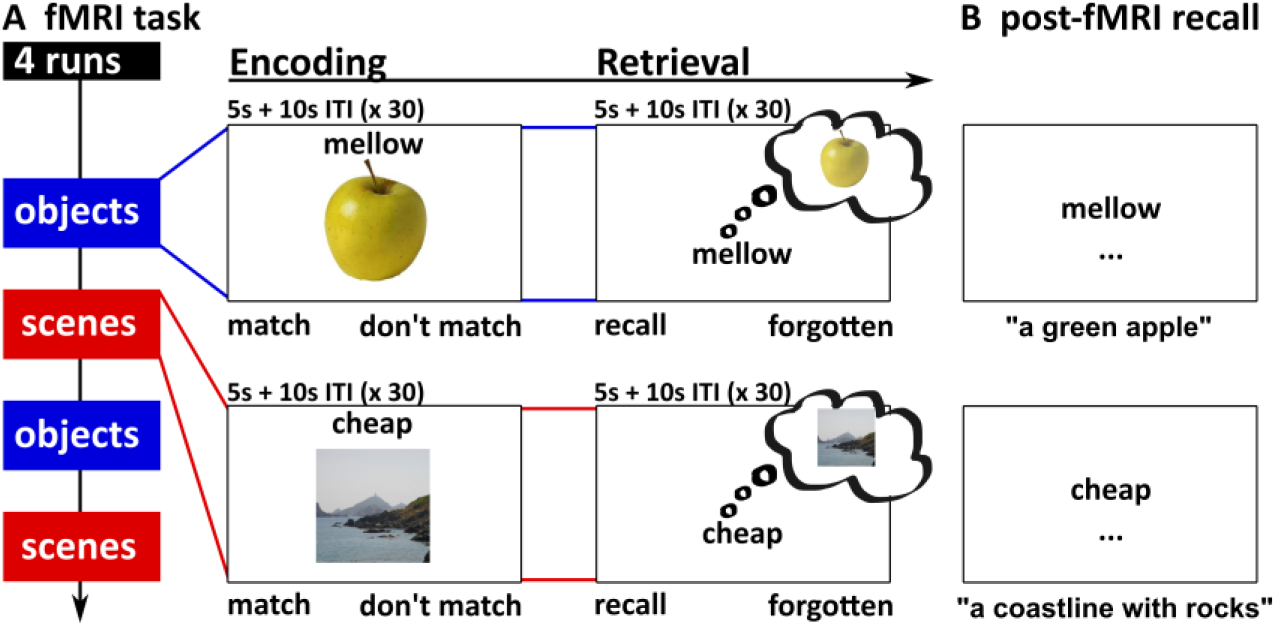
Experimental paradigm. **A**. The fMRI task consisted of two object and two scene runs, each comprising an encoding and a retrieval phase. During encoding, participants saw adjective-object or adjective-scene pairs. During retrieval, only the adjective was presented and participants tried to recall the associated object or scene from memory. Not shown: Each fMRI trial was followed by 10s of an active baseline task (ITI, arrows task), and the encoding phase was preceded and followed by a resting phase (odd-even numbers task, 180s) (see main text for details). **B**. In the post-fMRI recall task, participants typed in descriptions of the associated object and scene for each adjective.

Since memory responses given during the fMRI task were subjective, two measures were taken to ensure that the scanned retrieval portion accurately captured brain activity related to success vs. failure to recall. First, prior to the fMRI task, participants were explicitly instructed only to press ‘recall’ if they could vividly recall details of the associated image and to press ‘forgotten’ otherwise. Second, we additionally employed a post-fMRI recall task (Figure 1B) in order to obtain an objective memory measure. Again, participants were presented with each adjective, in the same order as during the fMRI retrieval phase. The task was to type a brief description of the associated image or a ‘?’ in case the target image was not recalled.

Critically, only trials with matching subjective and objective memory responses entered fMRI analyses (i.e. subjective ‘recall’ response during the fMRI task plus successful recall in the post-test, or subjective ‘forgotten’ response during the fMRI task plus unsuccessful recall in the post-test). This resulted in the following conditions of interest: object-recalled (OR), object-forgotten (OF), scene-recalled (SR), scene-forgotten (SF).

### 2.3 fMRI acquisition

Brain data were acquired using a GE Discovery MR750 3T system (GE Medical Systems) and a 32-channel head coil. For the functional runs, we used a gradient-echo, echo-planar pulse sequence (48 slices, 2.5mm isotropic voxels, TR=1000ms, TE=30ms, ascending acquisition order, multiband factor 3, 1300 volumes per run). The slice stack was oriented in parallel to the longitudinal MTL axis and covered nearly the whole brain (in some participants with larger brains, superior frontal cortex was not covered). The first 10 images of each run were discarded prior to analysis to allow for stabilization of the magnetic field. Additionally, a high-resolution whole-brain T1-weighted structural image (1×1×1mm, TR=7.9ms, TE=3.06ms) was acquired for each participant.

### 2.4 fMRI preprocessing and analysis

#### Regions of interest (ROI) strategy

Considering the high anatomical variability of the MTL (Pruessner et al., 2002), all analyses were carried out in unsmoothed, single-participant space within anatomical ROIs of the MTL (HC, PRC, PHC). These were hand-drawn on each participant’s T1 images using existing guidelines (Insausti et al., 1998; Pruessner et al., 2000, 2002), and resampled to functional space. To maximize object vs. scene sensitivity in the MTL cortex ROIs, considering gradual changes in content sensitivity along the parahippocampal gyrus (Litman et al., 2009; Liang et al., 2013), the posterior third of PRC and the anterior third of PHC were excluded from analysis (Staresina et al., 2011, 2012, 2013b). Across participants, the average number of voxels per bilateral ROI, in functional space and accounting for signal dropout, was 649.89 voxels (SEM: 15.07 voxels) for HC, 146.83 (11.68) for PRC, and 345.22 (10.92) for PHC.

#### Preprocessing

All analyses were carried out using Matlab and SPM12. Functional images were first corrected for differences in acquisition time (slice time correction), then corrected for head movement and movement-related magnetic field distortions using the ‘realign and unwarp’ algorithm implemented in SPM12. Structural images were then coregistered to the mean functional image before being segmented into grey matter, white matter, and CSF. Deformation fields from the segmentation procedure were used for MNI normalization (used for visualization only, see Figure 2A – all analyses were done in native space).

**Figure 2.**
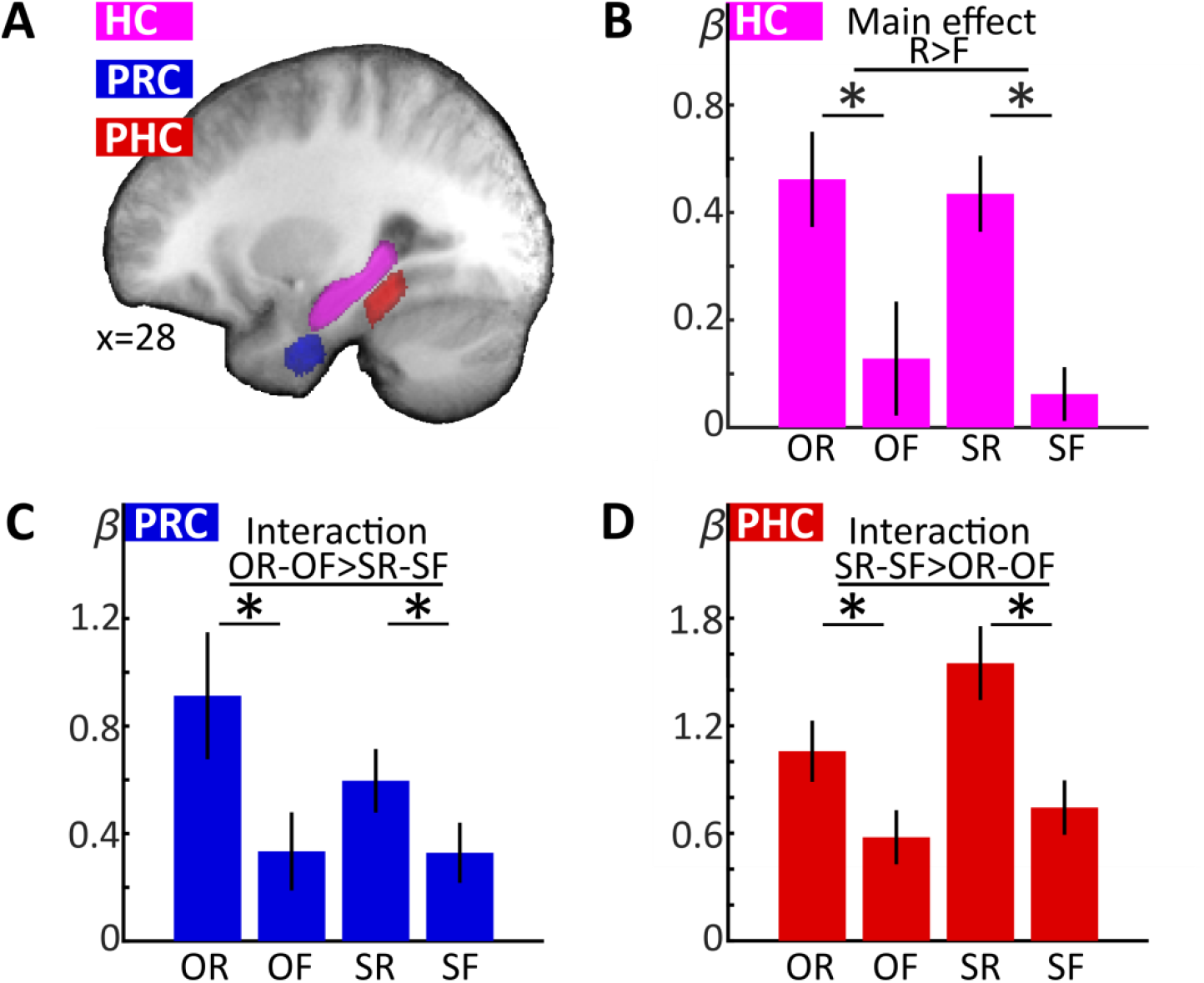
MTL ROIs and univariate retrieval results. **A**. To illustrate ROI localization, manually delineated ROIs for each participant’s HC, PRC, and PHC were MNI-normalized, averaged across participants and projected on the mean normalized T1 (averaged ROI threshold > 0.5). **B-D**. Retrieval-phase beta values were averaged within each participant’s individual ROIs and submitted to group analyses. HC (**B**) showed a main effect of successful recall, while PRC (**C**) and PHC (**D**) additionally showed interaction effects, indicating preference for object recall (PRC) and scene recall (PHC), respectively. O: object, S: scene, R: recalled, F: forgotten. Error bars denote SEM. \**p*<.05 (two-tailed) for pairwise *t*-tests.

#### Univariate analyses

For the first-level general linear model, all runs were concatenated and the high-pass filter (128s) and autoregressive model AR(1) + w were adapted to account for run concatenation. Regressors for our conditions of interest (OR, OF, SR, and SF for the encoding and retrieval phase, respectively) were modeled using a canonical hemodynamic response function (HRF) with a variable duration of each trial’s RT, assuming that memory-related processing of the stimulus is concluded at the time of the response. These regressors only included trials with matching memory responses during the fMRI task and post-fMRI recall. Non-matching trials (e.g. ‘recall’ response during the scan, but failed explicit recall during the post-scan) entered separate regressors of no interest. Additionally, the first-level model included non-convolved nuisance regressors for each volume of the transition and resting periods, and run constants. The resulting beta estimates from the retrieval phase were averaged across each participant’s ROIs before entering a group-level repeated-measures ANOVA with the factors region, content, and recall success. In case of sphericity violations, the degrees of freedom were adjusted using Greenhouse-Geisser correction.

#### Perception-retrieval overlap (PRO)

We asked whether, within each MTL cortex ROI (PRC, PHC), successful recall of a particular content is predicted, across voxels, by content tuning during perception. In that case, within PRC, there should be a positive correlation such that voxels that show stronger tuning to objects compared to scenes during perception should also be more engaged during successful compared to unsuccessful object recall. Similarly, within PHC, there should be a positive correlation such that voxels that show stronger tuning to scenes compared to objects during perception should be more engaged during successful compared to unsuccessful scene recall. This should be reflected in an across-voxel correlation of the effect sizes of the respective perception and recall contrasts, which we tested in the following way: We computed, for each participant, four *t* contrast images: (i-ii) the between-content perception contrasts from the encoding phase (O > S and S > O, irrespective of subsequent memory outcome); (iii) the within-content recall contrast for objects from the retrieval phase (OR > OF); (iv), the within-content recall contrast for scenes from the retrieval phase (SR > SF). *T* values across voxels were then vectorized for each participant and ROI. The perception-retrieval overlap for objects (PRO-O) was defined as the Pearson correlation coefficient between the object perception contrast *t* values (O > S) and the object recall contrast *t* values (OR > OF). Likewise, the perception-retrieval overlap for scenes (PRO-S) was defined as the Pearson correlation coefficient between the scenes perception contrast (S > O) and the scene recall contrast (SR > SF). Note that we only included voxels with positive values in the perception contrast (O > S for PRO-O, S > O for PRO-S) in this analysis to ensure that correlations are carried by voxels tuned to objects rather than scenes for PRO-O, and to scenes rather than objects for PRO-S. To ensure that these correlations would capture local rather than cross-hemispheric topographical relationships, the correlation coefficients were computed in left and right ROIs separately, then *Fisher z*-transformed and averaged. The resulting values were submitted to a two-way repeated measures ANOVA with the factors region (PRC, PHC) and correlation type (PRO-O, PRO-S), and followed up with two-sample and one-sample *t*-tests.

One possible concern is that PRO might be biased by temporal autocorrelations, which are greater within a run than between runs. Note though that the task consists of four functional runs, with two object- and two scene-only runs in alternating order. Each run contains an encoding and retrieval phase. Thus, in PRO, we correlate a contrast containing data from all four runs (O vs. S from all encoding phases) with contrasts containing data from only two runs (PRO-O: OR > OF; PRO-S: SR > SF). Consequently, both PRO-O and PRO-S correlate a contrast spanning all four runs with a contrast spanning two runs, making the overall temporal distance between contrasts equal. Moreover, whereas any bias arising from temporal autocorrelation would have similar impact across brain regions, we expect opposing patterns of PRO-O and PRO-S in PRC and PHC.

#### Control analysis 1: Specificity

In the above analysis, we correlate, across voxels of each ROI, the object perception contrast with the object recall contrast for PRO-O, and the scene perception contrast with the scene recall contrast for PRO-S. Importantly, we use only voxels with positive values in the perception contrast, i.e. object-selective voxels for PRO-O and scene-selective voxels for PRO-S. We expect positive values for PRO-O but not PRO-S in PRC, and for PRO-S but not PRO-O in PHC. However, one might argue that such results lack specificity: The object perception contrast in PRC may not only correlate with object recall (PRO-O), but also with scene recall. Similarly, the scene perception contrast in PHC may not only correlate with scene recall (PRO-S), but also object recall. This would indicate a non-specific relationship between perception and recall such that stronger content tuning during perception would predict stronger recall effects for either content. To control for this, we additionally computed the correlation between the object perception contrast (O > S, positive voxels only) and the scene recall contrast (SR > SF) for PRC, and the correlation between the scene perception contrast (S > O, positive voxels only) and the object recall contrast (OR > OF) for PHC.

#### Control analysis 2: Signal-to-noise ratio

Another possible concern might arise regarding the possible impact of differences in signal-to-noise ratio across voxels. Since the analysis is based on *t* contrasts between conditions, rather than estimates of activation in single conditions, we consider it unlikely that SNR gradients across voxels bias these results. Nevertheless, we additionally computed PRO as described above, but using partial Pearson correlations that included the temporal SNR of each voxel as a control variable. Temporal SNR was computed as the mean value of the preprocessed, unfiltered functional time series, divided by its standard deviation (separately per run, then averaged across runs).

## 3 Results

### 3.1 Behavioral results

We queried successful recall of objects and scenes at two time-points: During the fMRI task, participants merely responded ‘recall’ or ‘forgotten’ in response to each word cue (*subjective recall*). During a post-scan explicit word-cued recall task, participants typed in descriptions of the associated image, which were then scored by the authors (*objective recall*). Subjective responses during the fMRI task did not significantly differ by content (*t*_(17)_=0.685, *p*=.502), with nearly 50% ‘recall’ and ‘forgotten’ responses for both objects and scenes (mean [SEM] % ‘recall’ responses: objects: 51.2 [1.8], scenes 52.6 [2.6]). To test whether subjective ‘recall’ responses in the scanner were more likely to be followed by objective recall during the post-scan, we submitted the proportions of successful objective recall to a two-way repeated measures ANOVA with the factors content (objects, scenes) and subjective response (‘recall’, ‘forgotten’). This analysis yielded a significant effect of subjective response (*F*_(1,17)_=280.661, *p*<.001; no effect of content or interaction, *p*s≥.682); compared to subjective ‘forgotten’ responses, subjective ‘recall’ responses in the scanner were more likely to be followed by objective recall during the post-test for both objects (mean [SEM] % objective recall: 67.1 [4.3] after subjective ‘recall’ vs. 9.0 [1.6] after subjective ‘forgotten’) and scenes (66.5 [4.8] vs. 8.2 [2.2]). Note that only trials with consistent subjective and objective memory responses entered fMRI analysis (i.e. in the fMRI analysis, ‘R’ (‘recalled’) corresponds to a subjective ‘recall’ response during the fMRI task as well as accurate objective recall during the post-test; ‘F’ (‘forgotten’) corresponds to a subjective ‘forgotten’ response during the fMRI task as well as failed objective recall during the post-test). A repeated-measures ANOVA on the numbers of trials that entered fMRI analysis with the factors content (objects, scenes) and trial type (R, F) showed a significant effect of trial type (*F*_(1,17)_=6.473, *p*=.021), with more ‘F’ than ‘R’ trials (mean [SEM] number of trials: OR: 19.8 [1.5]. OF: 25.9 [1.3], SR: 20.1 [1.7], SF: 25.2 [1.6]; no effect of content or interaction, *p*s≥.668). All participants in the final sample contributed at least 8 trials per regressor of interest (OR, OF, SR, SF).

### 3.2 Content-independent vs. content-sensitive retrieval processing in MTL subregions

Univariate analyses were carried out within bilateral single-participant ROIs of HC, PRC, and PHC (Figure 2A). To characterize each MTL ROI with regard to its overall content-independent or content-sensitive response profile during retrieval, single-participant beta values for the regressors of interest (OR, OF, SR, SF) from the retrieval phase were averaged across all voxels for each individual’s ROI. ROI averages were then submitted to a repeated-measures three-way ANOVA with the factors region (HC, PRC, PHC), content (objects, scenes), and recall success (recalled, forgotten). This yielded a significant three-way interaction of region, content, and recall success (*F*_(1.46,24.80)_=10.014, *p*=.002), as well as significant two-way interactions of region with content (*F*_(1.42,24.10)_=13.544, *p*<.001) and region with recall success (*F*_(1.81,30.79)_=6.305, *p*=.006).

Subsequent analyses were carried out separately for each ROI, using two-way repeated-measures ANOVAs (including the factors content and recall success; Figure 2B-D). We expected content-independent recall in the HC, reflected in a main effect of successful recall. Conversely, we expected content-sensitive recall in the PRC and PHC, reflected in interaction effects of content and recall, with a preference for object recall in the PRC and scene recall in the PHC.

HC showed a significant main effect of successful recall (*F*_(1,17)_=24.509, *p*<.001), but no effect of content nor a recall success x content interaction (*p*≥.496). By contrast, PRC showed a significant main effect of successful recall (*F*_(1,17)_=18.137, *p*=.001), as well as a recall x content interaction (*F*_(1,17)_=4.579, *p*=.047) due to a stronger recall effect for objects relative to scenes. There was no main effect of content in PRC (*p*=.173). Finally, PHC showed a significant main effect of content (*F*_(1,17)_=16.804, *p*=.001), recall success (*F*_(1,17)_=27.329, *p*<.001), and a significant recall success x content interaction (*F*_(1,17)_=7.723, *p*=.013) due to a stronger recall effect for scenes relative to objects. To further characterize each ROI’s response profile, we computed post-hoc paired t-tests to assess object recall effects (OR vs. OF) and scene recall effects (SR vs. SF) in each ROI. All single comparisons were significant (*ts*_(17)_≥2.667, *ps*≤.016). Critically, however, as indicated by the above interaction effects, the object recall effect was greater than the scene recall effect in PRC, and vice versa in PHC. Taken together, the ROI results show content-independent recall-related activity in HC versus a preference for object recall activity in PRC and for scene recall activity in PHC.

### 3.3 Perception-retrieval overlap (PRO)

The preceding analysis established a preference for object recall in PRC and a preference for scene recall in PHC. Next, we assessed whether successful recall in these ROIs preferentially recruited voxels that were also diagnostic of object vs. scene perception during encoding. Note that this approach goes beyond a simple overlap of contrasts (as in a conjunction analysis): Rather than asking whether two contrasts exceed threshold in the same voxels, we ask whether there is a linear relationship between two contrasts such that voxels with a greater effect size in one contrast tend to show a greater effect size in the other (see Figure 3A for illustrative participant-level data). To this end, for each participant and ROI, we computed PRO-O (the correlation between the object perception contrast [O > S] from the encoding phase and the object recall contrast [OR > OF] from the retrieval phase), and PRO-S (the correlation between the scene perception contrast [S > O] from the encoding phase and the scene recall contrast [SR > SF] from the retrieval phase; see Methods for details). Note that PRO-O and PRO-S only included voxels tuned to either objects or scenes, as only voxels with positive values in the perception contrasts entered the correlation. We expected that PRC would show evidence for PRO-O: Voxels that are more tuned to objects over scenes during perception would be preferentially recruited during successful compared to unsuccessful object recall. In PHC, we expected evidence for PRO-S: Voxels that are more tuned to scenes over objects during perception would be preferentially recruited during successful compared to unsuccessful scene recall. We did not expect evidence for PRO-S in PRC or evidence for PRO-O in PHC.

**Figure 3.**
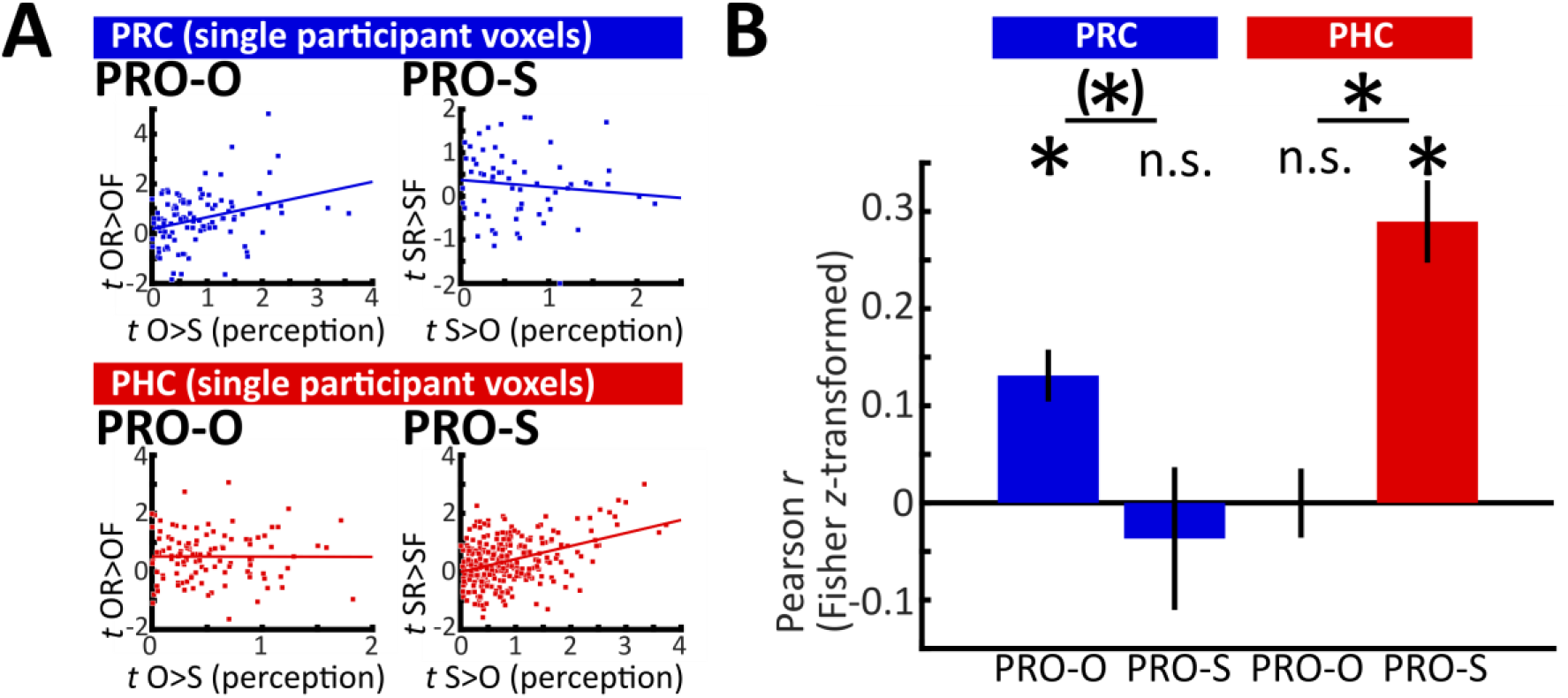
Perception-retrieval overlap (PRO). **A**. Illustrative data from two single participants’ ROIs. *T* values from the objects > scenes perception contrast (x axes, positive voxels only) are plotted against *t* values from the object recall contrast (PRO-O, left column), while *t* values from the scenes > objects perception contrast (positive voxels only) are plotted against the scene recall contrast (PRO-S, right column). Data points indicate single voxels. In these example data, PRC voxels with greater effect sizes for object perception tended to show greater effect sizes for successful object recall (upper left scatterplot). Similarly, PHC voxels with greater effect sizes for scene perception tended to show greater effect sizes for successful scene recall (lower right scatterplot). Note that these within-participant scatterplots are for visualization only. **B**. Group averages of *Fisher z*-transformed correlation coefficients for PRO-O and PRO-S for PRC and PHC. Across PRC voxels, object tuning predicted object recall (PRO-O), but scene tuning did not predict scene recall (PRO-S). Across PHC voxels, scene tuning predicted scene recall, but object tuning did not predict object recall. * *p*<.05, (*) *p*<.1 (two-tailed) for one-sample and paired *t*-tests, n.s.: not significant. Error bars denote SEM.

Before assessing the correlation between perception and retrieval contrasts, we confirmed PRC and PHC showed overall content tuning during perception. First, we tested whether the perception contrast yielded significant differences between objects and scenes when averaged across all voxels of each ROI. Second, we tested whether a majority of voxels in each ROI would show content tuning. Averaged across voxels, activation during object perception differed significantly from scene perception for both PRC (objects > scenes, *t*_17_=7.367, *p*<.001) and PHC (scenes > objects, *t*_17_=7.640, *p*<.001). As expected, HC showed no significant content tuning (*t*_17_=1.470, *p*=.160, numerically scenes > objects). Furthermore, the majority of PRC voxels showed object tuning, i.e. positive values in the O > S perception contrast (mean proportion: 63.70% [SEM: 1.60%]; one-sample t-test against 50%: *t*_17_=8.59, *p*<.001), whereas the majority of PHC voxels showed scene tuning, i.e. positive values in the S > O perception contrast (69.93% [2.04%], *t*_17_=9.78, *p*<.001). In the HC, the numerical majority of voxels were positive in the S>O contrast (S>O: 51.20% [1.16%]; *t*_17_=1.03, *p*=.315).

Results from the PRO analysis are summarized in Figure 3B. First, to confirm differences between PRC and PHC, we submitted the Fisher z-transformed correlation coefficients to a two-way repeated-measures ANOVA with the factors region (PRC, PHC) and content (PRO-O, PRO-S). This confirmed a significant interaction between region and content (*F*_1,17_=21.866, *p*<.001).

In PRC, correlation coefficients between O > S at encoding and the object recall contrast were significantly above zero (PRO-O, *t*_17_=4.910, *p*<.001), while correlation coefficients between S > O at encoding and the scene recall contrast were not (PRO-S, *t*_17_=0.500, *p*=.623). Furthermore, PRO-O trended to be greater than PRO-S (*t*_17_=2.073, *p*=.054). In contrast, in PHC, correlation coefficients between S > O at encoding and the scene recall contrast were significantly above zero (PRO-S, *t*_17_=6.832, *p*<.001), while correlation coefficients between O > S at encoding and the object recall contrast were not (PRO-O, *t*_17_=0.008, *p*=.994). PRO-S was significantly greater than PRO-O (*t*_17_=5.124, *p*<.001).

To test whether these findings are restricted to MTL cortical regions, we repeated the above analysis in HC. PRO-S, but not PRO-O, differed significantly from 0 (mean [SEM] PRO-O: 0.017 [0.030], *t*_17_= 0.565, *p*=.580, PRO-S: 0.059 [0.025], *t*_17_=2.330, *p*=.032). Furthermore, PRO-O and PRO-S did not differ from each other (*t*_17_=0.823, *p*=.422).

Our findings of PRO-O in PRC and PRO-S in PHC show that content tuning during perception in these ROIs predicts successful recall of that same content. To test the specificity of these findings, we repeated the analysis, this time testing whether content tuning would additionally predict recall of the non-preferred content. This would imply a non-specific relationship between content tuning during perception and recall. Hence, in PRC, we correlated the object perception contrast (O > S, positive voxels only) with the scene recall contrast. Correlation coefficients did not differ significantly from 0 (mean [SEM]: 0.031 [0.031], *t*_17_=1.011, *p*=.323), and were significantly smaller than PRO-O (*t*_17_=3.536, *p*=.003). In PHC, we correlated the scene perception contrast (S > O, positive voxels only) with the object recall contrast. Correlation coefficients were significantly greater than 0 (mean [SEM]: 0.120 [0.043], *t*_17_=2.789, *p*=0.013). Importantly, they were also significantly smaller than PRO-S (*t*_17_=5.007, *p*<.001). In sum, across PRC voxels, object tuning during perception predicted object recall (PRO-O) but not scene recall, and there was no relationship between scene tuning and scene recall. In contrast, across PHC voxels, scene tuning during perception predicted scene recall (PRO-S) to a greater extent than object recall, and there was no relationship between object tuning and object recall.

As a second control analysis, we computed PRO-O and PRO-S for PRC and PHC using partial Pearson correlations with each voxel’s temporal SNR as a control variable (see Methods). The statistical pattern was nearly identical for both the ANOVA and follow-up *t* tests, with the exception of the paired *t* test between PRO-O and PRO-S in PRC, which was now significant (*t*_17_=2.486, *p*=.024).

## 4 Discussion

Investigating cued recall of objects and scenes in the human MTL, we observed a triple dissociation across MTL subregions: While HC was engaged during successful recall of both content types, PRC preferentially tracked successful object recall and PHC preferentially tracked successful scene recall. Moreover, we demonstrate an across-voxel linear mapping of content-sensitive recall effects in PRC and PHC to content-tuning during the preceding encoding phase, suggesting that successful recall tends to draw on the same voxels that represent percepts with high specificity.

Before proceeding with the discussion, a note on terminology: Our results are agnostic to the debate whether MTL processing contributes to perception, or whether it necessarily serves a mnemonic function (Bussey and Saksida, 2007; Baxter, 2009; Suzuki, 2009, 2010; Graham et al., 2010; Squire and Wixted, 2011). We refer to the observed content tuning in the MTL cortex as perception as it results from sensory processing of objects and scenes, but we note that it may ultimately serve to encode representations into memory.

The present study provides strong evidence for an MTL memory model emphasizing an interplay of both content-sensitive and -independent modules: According to this view, PRC and PHC show differential involvement in object and scene processing, respectively, based on their anatomical connectivity profiles with the ventral and dorsal visual streams. HC links both circuits through direct and indirect (via entorhinal cortex) connections to PRC and PHC, implying a content-independent role of HC in memory (Suzuki and Amaral, 1994a, 1994b; Lavenex and Amaral, 2000; Davachi, 2006; Eichenbaum et al., 2007; van Strien et al., 2009; Wixted and Squire, 2011; Ranganath and Ritchey, 2012). Our findings of content-independent recall effects in HC, accompanied by preferential object recall in PRC and preferential scene recall in PHC, are in line with this view. Importantly, these connections are bidirectional (Suzuki and Amaral, 1994a, 1994b; Lavenex and Amaral, 2000; Eichenbaum et al., 2007), enabling information transfer from visual cortex via PRC/PHC to HC during perception and encoding, and vice versa during retrieval (Staresina et al., 2013b). This parallelism of MTL connectivity may underlie the phenomenon of cortical reinstatement - the reactivation of the same sensory cortical regions during recall that were already active during perception (Eichenbaum et al., 2007; Danker and Anderson, 2010). Our findings extend this concept: Even within content-sensitive cortical regions, voxels that are particularly tuned to one content type over the other during perception tend to be differentially reactivated when that content is successfully recalled. Importantly, such cortical reinstatement may underlie the psychological phenomenon of ‘re-living’ episodic memories during vivid recall (Eichenbaum et al., 2007; Danker and Anderson, 2010; Kuhl et al., 2011; Kuhl and Chun, 2014).

Our results constitute an important update to an existing body of work investigating content-sensitive recall. Previous studies have investigated cortical reinstatement by comparing cued retrieval of object-related and spatial information. However, most did not focus on differences between MTL cortices (Khader et al., 2005, 2007; Kuhl et al., 2011; Gordon et al., 2014; Kuhl and Chun, 2014; Morcom, 2014; Skinner et al., 2014; Bowen and Kensinger, 2017; Lee et al., 2018). Those that did are largely in line with our present findings. Staresina et al. (2012) showed that PHC reinstates scene information, while PRC reinstates low-level visual information (color). Similarly, Staresina et al. (2013b) demonstrated content-sensitive recall responses in PHC and PRC during retrieval of object-scene associations, driven by content-independent HC signals. One study presented evidence for object-cued reinstatement of faces in PRC, and object-cued reinstatement of places in PHC (Mack and Preston, 2016). However, that study only included correct memory responses, making it difficult to directly link the results to successful vs. unsuccessful recall. Finally, one study showed a dissociation between PRC vs. PHC for the reinstatement of an imagery task (person vs. place/object), but did not involve perceptual processing during encoding (Liang and Preston, 2017). Thus, the present study is the first to demonstrate a clear double dissociation between PRC and PHC during successful vs. unsuccessful object and scene recall triggered by a content-neutral cue, and to tie it to perceptual content tuning in a direct, voxel-wise manner.

Importantly, we observed preferential, but not exclusive, processing of objects and scenes in PRC and PHC, respectively. Both regions also show significant recall effects for their less-preferred content. Furthermore, in PHC, voxels that were tuned to scenes over objects during perception were also more active during successful object recall, albeit significantly less so than during scene recall. Previous studies have shown such overlap in content sensitivity in the MTL, with some object processing in PHC and some scene processing in PRC (Buffalo et al., 2006; Preston et al., 2010; Hannula et al., 2013; Liang et al., 2013; Martin et al., 2013; Staresina et al., 2013b; Martin et al., 2018). In particular, content sensitivity in the MTL cortex may not be abruptly demarcated, but follow a gradient (Litman et al., 2009; Liang et al., 2013). We sought to minimize this overlap by restricting analyses to the anterior two thirds of PRC and posterior two thirds of PHC, excluding the transition zone of the parahippocampal gyrus (Staresina et al., 2011, 2012, 2013b). Nevertheless, these two MTL subregions are not anatomically segregated, but show considerable interconnections (Suzuki and Amaral, 1994a; Lavenex and Amaral, 2000), facilitating cooperation. Furthermore, naturalistic scene images typically contain discernible objects, and many objects have a spatial/configurational component. A cardboard box, for example, may have the same general shape as a building, which has been shown to engage PHC (Epstein and Kanwisher, 1998). Similarly, object size modulates PHC activity (Cate et al., 2011; Konkle and Oliva, 2012). Thus, the significant (albeit weaker) responses of PRC and PHC during recall of their less-preferred content could stem from functional overlap in objects and scene processing, or from ambiguity in the stimuli themselves. Future studies could elucidate this ambiguity by controlling object and spatial features in these stimuli, albeit perhaps at the expense of decreasing natural validity.

It is important to note that our perception-retrieval overlap (PRO) differs from existing approaches that test for pattern similarity between encoding and retrieval (pattern reinstatement). For instance, ‘encoding-retrieval similarity’ (ERS) has shown that, relative to forgotten trials, successfully remembered trials are more similar to their respective encoding trials (Staresina et al., 2012; Ritchey et al., 2013). Similarly, in multivariate pattern analysis (MVPA), a classifier may be trained on encoding trials to distinguish between voxel patterns associated with different content or tasks, and then tested on retrieval trials (e.g. Polyn et al., 2005; Johnson et al., 2009; Mack and Preston, 2016; Liang and Preston, 2017). PRO, on the other hand, relies on across-voxel correlations of contrasts, rather than single conditions or trials. Previously, Haxby and colleagues used contrast correlations (Haxby et al., 2001) to demonstrate that content tuning during perception is stable between runs. Here, we test contrast correlations between tasks; specifically, whether content tuning – i.e. the difference between object and scene responses – can predict activity associated with successful recall of objects (PRO-O) and scenes (PRO-S) across voxels. Thus, PRO could be considered a more constrained form of a conjunction, or inclusive masking, analysis. These methods test whether two or more contrasts exceed some threshold in the same voxels - implying topographical overlap of the constituting contrasts, but, critically, not a positive correlation across voxels. While our results likely reflect cortical reinstatement, they illuminate a distinct aspect of it compared to pattern similarity in the sense of ERS and MVPA: The latter methods demonstrate that distributed patterns of activity associated with a certain content are reinstated during recall, while our results link content-sensitive recall effects to voxels that are highly tuned to that content over another. A similar link has been demonstrated between content tuning and recognition memory for PRC activity (Martin et al., 2016): In that study, distributed voxel patterns in PRC that were diagnostic of face recognition also showed face-sensitive perceptual tuning. However, that study did not establish a linear relationship.

Taken together, our results support an MTL model of episodic memory based on anatomical connectivity and demonstrate a direct topographical mapping between content-sensitive perception and recall in the MTL cortex. One remaining question is how mnemonic content is conveyed and transformed from content-sensitive MTL cortex to content-independent HC. Much of the information exchange between PRC/PHC and HC is relayed via the entorhinal cortex and its anterolateral and posteriormedial subregions (Suzuki and Amaral, 1994b; Maass et al., 2015; Navarro Schröder et al., 2015), which have similarly been shown to support content-sensitive processing (Schultz et al., 2012; Reagh and Yassa, 2014; Navarro Schröder et al., 2015; Berron et al., 2018), albeit potentially in a more integrated fashion (Schultz et al., 2015). How entorhinal retrieval processing relates to content tuning is unclear, though there is evidence for reinstatement of encoding representations in the entorhinal cortex (Staresina et al., 2013a). Future research may investigate the relationship between encoding and retrieval in the entorhinal cortex by making use of advanced high-resolution and ultra-high field approaches, thereby enhancing our understanding of the human MTL in its entirety.

## Acknowledgments

This work was supported by a European Commission Marie Skłodowska-Curie Fellowship (752557) to H.S., a Wellcome Trust/Royal Society Sir Henry Dale Fellowship (107672/Z/15/Z) to B.P.S., and a grant from the Marcus and Amalia Wallenberg Foundation to A.D.W.

